# Single unit activity in marmoset posterior parietal cortex in a gap saccade task

**DOI:** 10.1101/737312

**Authors:** Liya Ma, Janahan Selvanayagam, Maryam Ghahremani, Lauren K. Hayrynen, Kevin D. Johnston, Stefan Everling

## Abstract

Abnormal saccadic eye movements can serve as biomarkers for patients with several neuropsychiatric disorders. To investigate cortical control mechanisms of saccadic responses, the common marmoset (*Callithrix jacchus*) is a promising non-human primate model. Their lissencephalic brain allows for accurate targeting of homologues of sulcal areas in the macaque brain. Here we recorded single unit activity in the posterior parietal cortex of two marmosets using chronic microelectrode arrays, while the monkeys performed a saccadic task with Gap trials (stimulus onset lagged fixation point offset by 200ms) interleaved with Step trials (fixation point disappeared when the peripheral stimulus appeared). Both marmosets showed a gap effect—shorter saccadic reaction times (SRTs) in Gap vs. Step trials. On average, stronger gap-period response across the entire neuronal population preceded shorter SRTs on trials with contralateral targets, although this correlation was stronger among the 15% ‘gap neurons’, which responded significantly during the gap. We also found 39% ‘target neurons’ with significant visual target-related responses, which were stronger in Gap trials and correlated with the SRTs better than the remaining cells. Compared with slow saccades, fast saccades were preceded by both stronger gap-related and target-related response in all PPC neurons, regardless of whether such response reached significance. Our findings suggest that the PPC in the marmoset contains an area that is involved in the modulation of saccadic preparation and plays roles comparable to those of area LIP in macaque monkeys in eye movements.

**SIGNIFICANCE STATEMENT:** Abnormal saccadic eye movements can serve as biomarkers for different neuropsychiatric disorders. So far, processes of cerebral cortical control of saccades are not fully understood. Non-human primates are ideal models for studying such processes, and the marmoset is especially advantageous since their smooth cortex permits laminar analyses of cortical microcircuits. Using electrode arrays implanted in the posterior parietal cortex of marmosets, we found neurons responsive to key periods of a saccadic task in a manner that contribute to cortical modulation of saccadic preparation. Notably, this signal was correlated with subsequent saccadic reaction times and was present in the entire neuronal population. We suggest that the marmoset model will shed new light on the cortical mechanisms of saccadic control.

## INTRODUCTION

Saccades are ballistic, conjugate eye movements that sample the visual environment and thereby serve as a gateway to higher cognitive functions. Distinct deficits in saccadic tasks can serve as both diagnostics for various neuropsychiatric disorders and indicators for underlying changes in the oculomotor network (Klein et al., 2000; Munoz and Everling, 2004; Hutton and Ettinger, 2006; Gooding and Basso, 2008). While the brainstem and superior colliculus (SC) control the generation of saccades (Waitzman et al., 1991; Keller and Edelman, 1994), attention and other cognitive processes strongly modulate the reaction times of these movements (Hutton, 2008). Although known to be mediated by cortical areas including the frontal eye fields (FEF) and posterior parietal cortex (PPC) (Gaymard et al., 1998; DeSouza et al., 2003; Brown et al., 2004), the detailed microcircuits and computations involved in these processes are still not fully understood.

The common marmoset is a burgeoning primate model that shares homologous functional networks with humans and macaque monkeys (Ghahremani et al., 2017)—the most common non-human primate model. Behaviorally, the marmoset holds enormous potential for the study of primate communication and social behaviors (Miller et al., 2016). Given that eye movements are a crucial gauge for cognitive processes in primate studies, understanding the cortical mechanisms of oculomotor control is fundamental to cognitive studies in marmosets. Recent studies have demonstrated that the marmosets can be trained to perform oculomotor tasks (Johnston et al., 2018) and display visual behaviors comparable to the macaques (Mitchell et al., 2014, 2015). Anatomically, the marmoset has a smooth cerebral cortex (lissencephaly) which permits laminar analyses of local microcircuits in areas that are hidden in sulci in the macaque monkey, such as the FEF and those in the banks of the intraparietal sulcus (IPS). So far, both anatomical (Collins et al., 2005; Reser et al., 2013) and resting-state functional MRI studies (Ghahremani et al., 2017) have identified a putative homologue of the macaque lateral intraparietal area (LIP) in the marmoset PPC, based on its connections with both SC and FEF. A recent study demonstrated that microstimulations in the marmoset PPC evoked eye blinks and saccades (Ghahremani et al., 2019), as can be expected from an area homologous to area LIP in the macaques (Shibutani et al., 1984; Kurylo and Skavenski, 1991; Thier and Andersen, 1996, 1998). To date, however, no single-unit recording study has been conducted to characterize the involvement of the marmoset PPC neurons in saccadic tasks.

In saccade tasks conducted in both humans and macaques, a brief ‘gap’ (typically 200ms) intervening between the offset of the fixation point and the onset of the peripheral target is known to shorten subsequent saccadic reaction times (SRTs) (Saslow, 1967) and elicit saccades with very short latencies— so called express saccades (Fischer and Boch, 1983). In macaque monkeys, increased activity in the gap period and higher pre-target activity levels for express compared with regular saccades were found in individual neurons in the SC (Dorris et al., 1997; Everling et al., 1999), the FEF (Dias and Bruce, 1994; Everling and Munoz, 2000) as well as in area LIP (Chen et al., 2013, 2016). In humans, EEG signals from the occipital-parietal network became enhanced before express saccades (Everling et al., 1996). The reduction in SRTs afforded by the gap was attributed to both fixation release (Fendrich et al., 1991; Reuter-Lorenz et al., 1991; Sommer, 1994; Dorris and Munoz, 1995), and advanced preparation of saccadic motor programs (Paré and Munoz, 1996). In comparison, the generation of express saccades can be explained by motor preparation alone as demonstrated in the SC (Dorris et al., 1997; Everling et al., 1998), the FEF (Everling and Munoz, 2000) and area LIP in macaque monkeys (Chen et al., 2013, 2016). Given that the marmoset PPC directly projects to the SC (Collins et al., 2005) and is reciprocally connected with frontal areas which may correspond to the marmoset FEF (Reser et al., 2013; Majka et al., 2016), we hypothesize that it contains an area that plays a similar role in the motor preparation preceding the gap effect and express-like saccades as area LIP in macaques.

Here we report results from microelectrode array recordings from the PPC of marmoset monkeys while they performed visually-guided saccades with or without a 200-ms gap period (Gap vs Step trials) in a randomly interleaved fashion. As we showed previously (Johnston et al. 2018), marmosets displayed shorter SRTs in the Gap than Step trials. Of the sample of recorded single PPC neurons, 15% exhibited significant modulations of activity during the gap, which we will refer to as ‘gap neurons’. Across all neurons, stronger gap-period responses preceded shorter SRTs on trials with subsequent contralateral targets, although such negative correlation was more prominent among gap neurons. Additionally, we found that 39% of PPC neurons responded to the saccadic target with an enhanced response in Gap trials, which we will refer to as ‘target neurons’. Their responses also negatively correlated with subsequent SRTs. As expected from these negative correlations, both types of neurons showed stronger responses before fast saccades with SRTs in the shortest quartile, compared to the rest of the distribution which we called ‘slow saccades’. Interestingly, the gap-related response of the remaining 85% non-gap neurons was on average still significantly greater on trials with fast than those with slow contralateral saccades, and so was the target-related response of the 61% non-target neurons, suggesting a widespread performance-related signal. Our findings suggest that the marmoset PPC contains an area that plays a similar role as the macaque LIP in modulating saccadic motor preparation, which contributes to the gap effect and the generation of fast saccades in the gap task.

## MATERIALS AND METHODS

### Animals

Two male common marmosets (*Callithrix jacchus*), weighing 440 g and 451g at the age of 2.5 and 4yrs respectively were used in the study. All procedures performed were approved by the Animal Care Committee of the University of Western Ontario Council on Animal Care and in accordance with the Canadian Council of Animal Care policy on laboratory animal use.

After the initial acclimatization to the custom-designed chair restraint, over the course of several weeks, the marmosets were gradually trained to sit quietly facing the monitor and to consistently lick the sipper tube for their preferred liquid reward, which was delivered intermittently (Johnston et al., 2018). For Marmoset B, the preferred reward was sweetened condensed milk mixed with water in a 2:1 ratio, and for Marmoset W this was corn syrup mixed with water in a 1:1 ratio. Once they could sit calmly for 45 minutes, the first surgery was performed to install a head restraint/recording chamber (Johnston et al., 2018). A second microelectrode array implantation surgery took place after the monkeys became proficient with the behavioral task.

### Surgical procedures

For both surgeries, the animals were anesthetized with ketamine and maintained with intravenous propofol and gaseous isoflurane (Johnston et al., 2018). Their heart rate, SpO2, temperature and breathing were continuously monitored by an experienced veterinary technician. After each surgery, the marmosets received postsurgical treatments including analgesics and antibiotics to minimize pain or discomfort, under the oversight of a university veterinarian.

In the first surgery, a custom-designed combination head restraint/recording chamber (Johnston et al., 2018) was attached to the skull with UV-cured dental adhesive and resin (All-Bond Universal and Duo-Link, Bisco Dental Products (Canada), Richmond, BC, Canada). Together with a custom-designed protective cap, the chamber would serve to protect the electrode array after its implantation. After the animals were well-trained on the task, they underwent a second surgery, in which a parietal craniotomy was made inside the recording chamber at 1.4mm anterior and 6mm lateral to the interaural midpoint and a 32-channel Utah array (Blackrock Microsystems, Salt Lake City, UT, USA) was implanted. The positioning of the array was guided by both stereotaxic coordinates of area LIP (Paxinos et al., 2012) and the location of a posterior parietal area functionally connected to the SC (Ghahremani et al., 2017). Additionally, we were guided by the location of a small blood vessel which corresponded to the location of the shallow IPS. Before insertion of the array, we secured a ground screw in a small burr hole made posterior to the craniotomy. Arrays were manually inserted so that they straddled the IPS and covered as much of the sulcus as possible along the anterior-posterior axis. The connecting wires and the connector were secured inside the chamber using dental resin. The grounding wires were then tightly wound around the ground screw to ensure electrical connection before being secured with dental resin. The array and craniotomy were protected by a very fine layer of gel foam and medical-grade silicone elastomer adhesive (Kwik-Sil, World Precision Instruments, Sarasota, FL, USA) before being covered by dental resin.

### Behavioral training and paradigm

On the day before each training or recording session, the animals received mild food restriction: the size of their second of two daily meals was reduced to 80% of their ad libitum consumption amount. On the training/recording day, the session always took place before the first of their two daily meals (Johnston et al., 2018). During training, the animals received their preferred liquid reward (for details see previous section) upon the successful completion of each trial.

Training on the goal-directed saccade task consisted of two steps: fixation training and saccade training, which were described in detail previously (Johnston et al., 2018) and summarized briefly here. During fixation training, the animals learned to start fixating within 4s and maintain the gaze for 500ms on a marmoset face 0.8° × 0.8° in size within a 5° × 5° electronic window. The possible location for this stimulus was gradually increased from 1 (centre only) to 3 (centre, left and right) and was used in a random order. Once the animal was able to respond to the stimulus in each location as required, the electronic window was reduced to 3° × 3°. During saccade training, if the fixation was maintained successfully for 500ms, the central stimulus was replaced immediately by a second stimulus presented at 5° to the left of fixation. The animals were rewarded if they initiated a saccade to the target stimulus within 1s and maintained fixation for 10ms on the target. Once the animal acquired this response, a second possible target 5° to the right of fixation was used, first on alternating trial blocks with the left target, then on randomly selected trials. When the animals became proficient at the task, we reduced the time allowed between fixation stimulus offset and saccade onset to 500ms and replaced each face stimulus with a white dot (0.25° in diameter, luminance 10 cd/m^2^) at the centre of the 5° × 5° window. The same fixation dot was then used in the goal-directed saccade paradigm. The same dark background (2 cd/m^2^) was used in both training and the goal-directed saccade task.

The goal-directed saccade paradigm included Step and Gap conditions (Figure 1A). Each trial began with the appearance of a white dot (see above for details) at the centre of the screen. The animals were required to initiate fixation within 3s and maintain the gaze within a window of 1.5° × 1.5° for 700-900ms. On Step trials, concurrent with the offset of the fixation spot, a peripheral target was presented pseudorandomly to the left or right by 5°. The target is a white dot greater in size (0.8° in diameter) and equiluminesecent with the fixation dot (10 cd/m^2^). The animals were rewarded if they generated a saccade within 500ms and if the saccade endpoint fell within a 3° × 3° window surrounding the target. On Gap trials, after 500-700ms of fixation, the spot was extinguished for a ‘gap period’ of 200ms, during which the animals were required to maintain their gaze within the same electronic window. The onset of a peripheral target marked the end of the gap period. The animals were rewarded if they generated responses that met the same criteria as in the Step condition. Gap and Step trials were randomly interleaved. The animals’ eye positions were recorded and digitized at 1000Hz using an Eyelink 1000 infrared pupillary tracking system (SR Research, Mississauga, ON, Canada).

**Figure 1.**
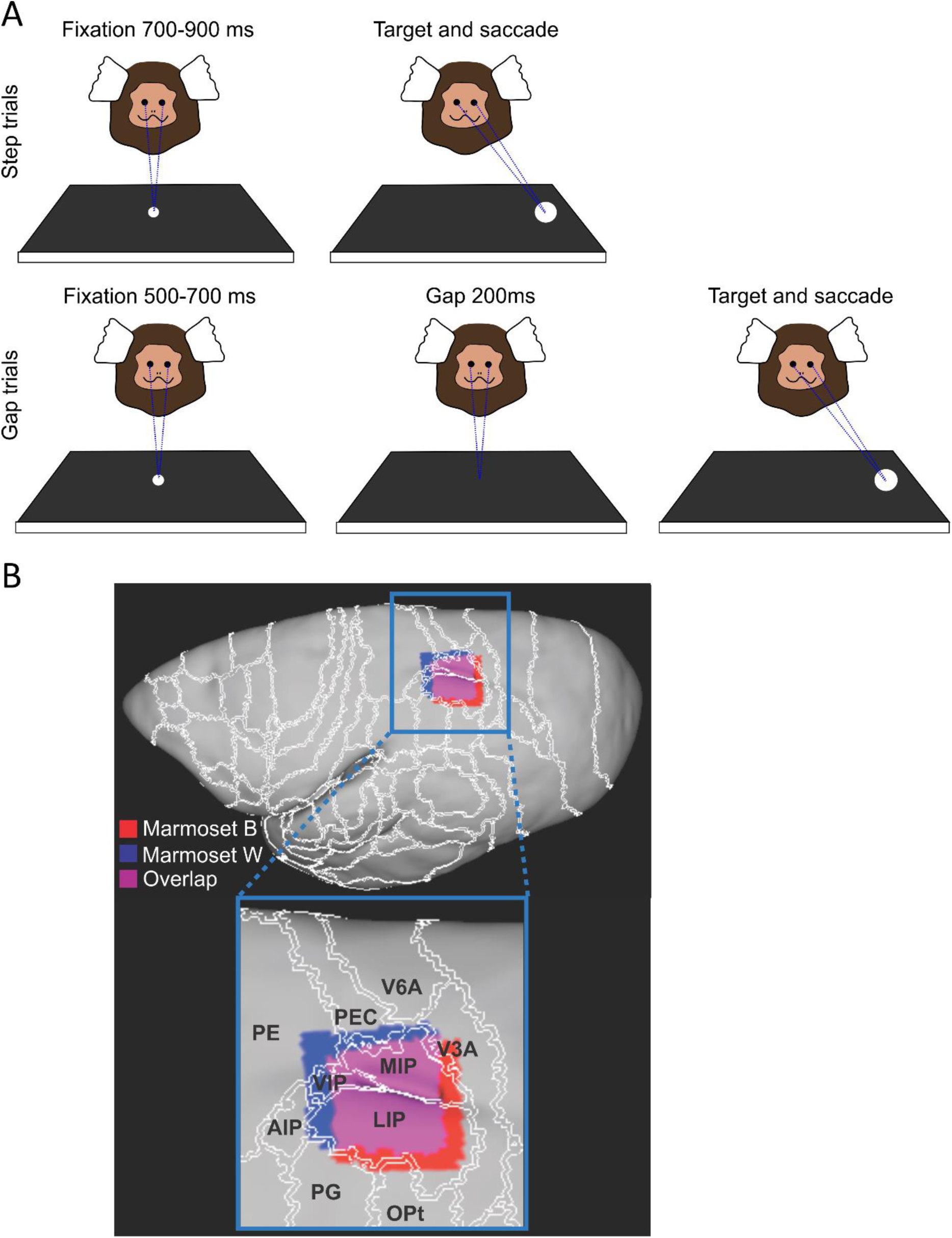
Schematic illustration of the goal-directed saccade paradigm and positioning of the microelectrode arrays. A). The two trial types and the left/right location of the saccadic target were randomly interleaved. In Step trials, the animals had to maintain fixation on the central white dot for a random interval between 700 and 900ms before its offset and the onset of the peripheral target, to which they had to make a saccade (top row). In Gap trials, the fixation dot extinguished after an interval between 500 and 700ms, and the target came on after a 200-ms gap period during which fixation had to be maintained despite of the lack of any visual display (bottom row). A liquid reward was delivered if the animal made a saccade to the target within 500ms. B). Array positioning based on ex vivo MRI and in vivo micro-CT scan for the two marmosets, respectively. Top: array locations for Marmoset B (red) and Marmoset W (blue) registered on the surface space of the left hemisphere of the brain, with their overlap shown in purple and cortical boundaries overlaid in white. Bottom: zoomed-in view of the recorded area with neighboring areas labelled according to the NIH marmoset brain parcellation map (Lui et al 2018). LIP: lateral intraparietal area, MIP: medial intraparietal area, VIP: ventral intraparietal area, AIP: anterior intraparietal area, PE: parietal area PE, PEC: caudal part of the parietal area PE, PG: parietal area PG, OPt: occipito-parietal transition area, V6A: visual area 6A, V3A: visual area 3A.

### Recording and data analysis

Neural activities, including LFP and spike trains, as well as eye-tracking data were recorded using a multiacquisition processor (MAP) system (Plexon, Dallas, TX, USA) for Marmoset B and the Open Ephys acquisition board (http://www.open-ephys.org) and digital headstages (INTAN, Los Angeles, CA) in Marmoset W. Data collected with both systems were converted to Neuroexplorer (nex) files and single units were isolated by applying principal component analysis in 2D and 3D with the Plexon Offline sorter (Plexon, Dallas, Texas) and analyzed in MATLAB (MathWorks, Naticks, MA, USA, RRID:SCR_001622). Single units with firing rates less than 0.3Hz were excluded from further analyses.

For behavioral analysis, we included only correctly performed trials with saccadic reaction times (SRTs) less than 350ms. We also identified anticipatory saccades—responses initiated before the visual target was processed—by plotting the cumulative percentage of correct responses as a function of SRT (Figure 2). For each trial type and in each animal, we found a point of clear change in slope. At and above these points, the performance of the animals exceeded 70% in each trial type (Figure 2). We therefore used these turning points as empirical cut-offs and trials with SRTs that fell below these cut-off SRTs were excluded from further analyses as anticipatory saccades.

**Figure 2.**
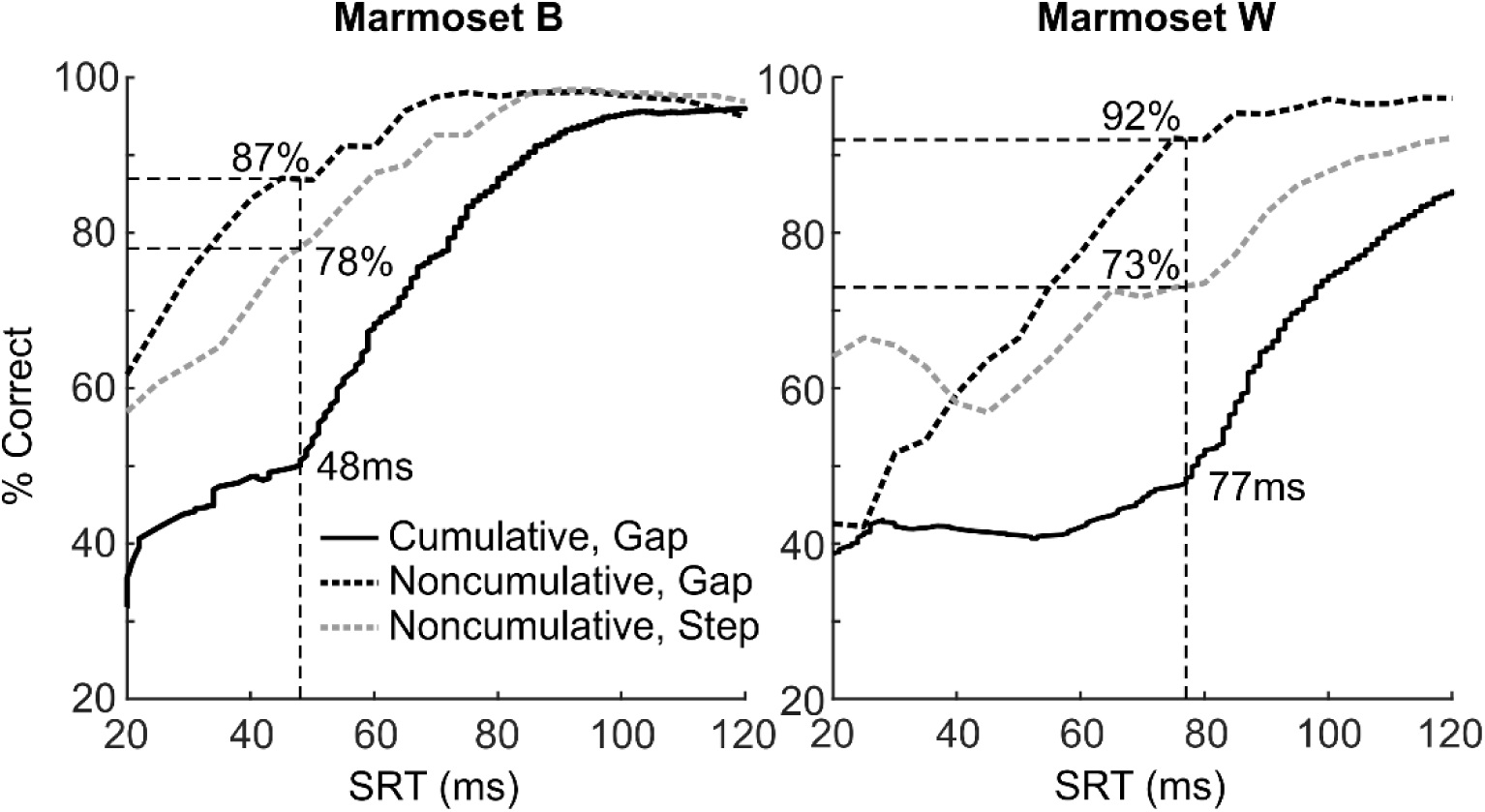
Percentages of correct saccadic responses plotted as a function of SRTs. The sudden change in the slope (vertical dashed line) of the cumulative percentage of correct responses (black solid curve) is used as the empirical threshold for anticipatory saccades. At this threshold, the animals’ noncumulative performance, i.e. percentage correct in consecutive time bins, was above 70% in both Gap (black dashed curve) and Step (gray dashed curve) trials. Left: performance as a function of SRT in Marmoset B. Right: performance in Marmoset W. SRT, saccadic reaction time.

For the analyses of neuronal activities, we focused on three behavioral epochs: the fixation period, the peristimulus period and the visual period. The fixation period was defined as the interval from 400ms to 201ms before the onset of the peripheral target. In Gap but not Step trials, the end of the fixation period coincided with the offset of the fixation point, or the onset of the gap period. The peristimulus period was defined as the interval from 165ms before to 34ms after the onset of the peripheral target. In Gap trials, this period nearly coincided with the gap period, starting 35ms after the offset of the fixation dot (Figure 1A). The first 34ms after target onset still belonged to the gap period because that single-unit target-related response started no sooner than 35ms after target onset (Figure 6C). This may suggest that the visual signal takes at least 35ms to reach the PPC in the marmosets. The visual period was defined as the interval from 35ms to 134ms after target onset, 100ms in duration. Given that it ended later than the reaction times of many saccades, activities captured in this window could also reflect post-saccade processes.

Parametric statistical tests such as *t-*tests and ANOVAs were used provided that each group involved in the test had a sample size greater than 30. For the correlation analysis (Figure 8) we used the nonparametric Spearman’s rho, because the parametric Pearson’s rho assumes strict normality in each group of data tested. To determine whether the correlation coefficients were significantly different from zero at the group level, we used one-sample *t-*tests with adjustments for family-wise false discovery rate (Benjamini and Hochberg, 1995; Groppe et al., 2011). Nonparametric Komolgorov-Smirnov test was used to determine whether a type of neurons were distributed similarly across animals, or whether different types of neurons were similarly distributed.

### Spike Density Function

To evaluate the relationship between neural activity and stimulus onset and saccade onset, continuous spike density functions were constructed. The activation waveform was obtained by convolving each spike with an asymmetric function that resembled a postsynaptic potential (Hanes and Schall, 1996; Thompson et al., 1996). The advantage of this function over a standard Gaussian function (Richmond and Optican, 1987) is that it accounts for the fact that spikes exert an effect forward, but not backward in time. We used this convolution method in all line plots of single neuron or group averaged activity, but not on data used in any statistical tests or shown in bar graphs.

### Confirming Array Location

Ex-vivo MRI was conducted to confirm the positioning of the array for Marmoset B. As Marmoset W is involved in additional experiments, in vivo micro-CT scan was used as an alternative method to confirm the array location.

#### Ex vivo MRI scan

To prepare for the MRI, Marmoset B was euthanized through transcardial perfusion and its brain was extracted at the end of the procedure. Anesthesia was induced with 20 mg/kg of ketamine plus 0.025 mg/kg Medetomidine and maintained with 5% isoflurane in 1.4-2% oxygen at a state deeper than the surgical plane, with no response to cornea touching or toe pinching. The animal was then transcardially perfused with 200 ml of phosphate buffered saline, followed by 200 ml of 10% formaldehyde buffered solution (formalin). The brain was then extracted and stored in 10% buffered formalin for over a week. On the day of the scan, the brain was transferred and immersed in a fluorine-based lubricant (Christo-lube, Lubrication Technology, Inc) to improve homogeneity and avoid susceptibility artifacts at the boundaries. The ex-vivo image was then acquired using a 9.4T, 31 cm horizontal bore magnet (Varian/Agilent) and Bruker BioSpec Avance III console with the software package Paravision-6 (Bruker BioSpin) and a custom-built 15-cm-diameter gradient coil with 400 mT/m maximum gradient strength (xMR, London, Ontario, Canada; Peterson et al., 2018). An ex-vivo T2-weighted image was acquired with the following scanning parameters: repetition time (TR) = 5s, echo time (TE) = 45 ms, field of view (FOV) = 40 × 32, image size = 160 × 128, slice thickness = 0.5 mm.

To identify the location of the array, the resulting T2-weighted image was registered to the NIH marmoset brain atlas (Liu et al., 2018) using the registration packages of the FSL software (fMRI Software Library: http://www.fmrib.ox.ac.uk). Upon visual examination of the image, an indentation of comparable size to the array (2.4 × 2.4 mm) was identified on the surface of the cortex within the PPC that represented the array location. The location of this region of interest was interpolated on the cortical surface to create a mask across this indentation, meant to represent an approximation of the array location. The mask was then projected onto the surface space in CARET toolbox (Van Essen et al., 2001), using a surface-based version of the NIH volume template that was kindly provided by the authors of the NIH marmoset brain template (Liu et al., 2018). The array mask was then compared to the area LIP as defined by the parcellated regions of the NIH template, that was also projected on CARET surface space.

#### In vivo micro-CT scan

Marmoset W was imaged in a live-animal micro-CT scanner (eXplore Locus Ultra, GR Healthcare Biosciences, London, ON) to identify the array location. Prior to the scan, the animal was anesthetized with 15mg/kg Ketamine mixed with 0.025mg/kg Medetomidine. He was then placed on his back on the CT bed with arms positioned down along his sides and then inserted inside the scanner. X-ray tube potential of 120 kV and tube current of 20 mA were used for the scan, with the data acquired at 0.5° angular increment over 360°, resulting in 1000 views. The resulting CT images were then reconstructed into 3D with isotropic voxel size of 0.154 mm. Heart rate and SpO2 were monitored throughout the session. At the end of the scan, the injectable anesthetic was reversed with an IM injection of 0.025mg/kg Ceptor.

The location of the array was clearly identified within marmoset PPC by visual inspection of the CT image. To find the location of the array with respect to the NIH template, the acquired CT image was brain extracted while including the trace of the array across the boundary of the cortex. The brain extracted image was then registered to the NIH marmoset brain atlas (Liu et al., 2018) using the FSL software (fMRI Software Library: http://www.fmrib.ox.ac.uk). Similar to the ex-vivo MRI data, an ROI mask was created over the traces of the array across the surface of the cortex to represent the location of the array. This mask along with the actual location of area LIP was projected on the surface space using CARET, to compare the positioning of the array to area LIP.

## RESULTS

We recorded neural activity through 32-channel Utah arrays in 13 sessions in Marmoset B and 14 sessions in Marmoset W, while the monkeys performed randomly interleaved Gap and Step trials (Figure 1A). The recording locations in the posterior parietal cortex (PPC), identified using either in vivo micro-CT scan or ex-vivo MRI combined with the NIH marmoset brain atlas (Liu et al., 2018), are shown in Figure 1B.

### Behavior

In total, Marmoset B performed 2662 correct trials and Marmoset W performed 2729 correct trials. The sessions had a mean duration of 32.6 min (s.d. = 8.2min). Trials where the animal made saccades that were anticipatory, incorrect or with saccadic reaction times (SRTs) greater than 350ms were excluded from further analyses. To define anticipatory saccades empirically, we plotted the cumulative percentage of correct saccadic responses with SRTs equal to or less than each given SRT threshold for each subject (Figure 2). In each plot, the abrupt increase in the slope of the solid line marks the saccades with the shortest SRTs that were informed by the actual target location, rather than being generated anticipatorily (dashed lines with colors matching the curve; Figure 2). At these empirical cut-offs, the animals’ performance on both trial types (dashed curves) were greater than 70%. In the following analyses, we included only correct trials with SRTs greater than 48ms and 77ms respectively for Marmoset B and W.

For trials with responses that met the criteria, we plotted the distributions of SRTs in Gap and Step trials, respectively (Figure 3A and B). The black bars indicate the percentage of correct trials in each SRT bin, the gray bars denote the percentage of error trials and the empty bars show the percentage of anticipatory saccades. We also separated the SRTs by saccadic direction: ipsilateral saccades towards visual targets on the same side as the electrode arrays (bottom panels in Figure 3A and B) and contralateral saccades towards the opposite side (top panels in Figure 3A and B). All these distributions were significantly non-normal in both animals (Komolgorov-Smirnov tests, KS stats >= 0.5, p <= 5.0 × 10^−20^), thus non-parametric tests were used to compare SRTs across tasks. In both marmosets, the SRTs of correct trials were significantly shorter during Gap than Step trials (rank sum test, Marmoset B: Z = –6.32, p = 2.6 × 10^−10^, Marmoset W: Z = –13.4, p = 3.9 × 10^−41^). Unlike macaque monkeys, but not unlike humans, the SRTs in Gap trials were not distinctively bimodally distributed. Hence, we could not objectively separate the responses into express and regular saccades. Instead, we separate the trials into those with fast and slow saccades, based on whether their SRTs fell into the shortest quartile or not (Figure 3A and B, red dashed lines).

**Figure 3.**
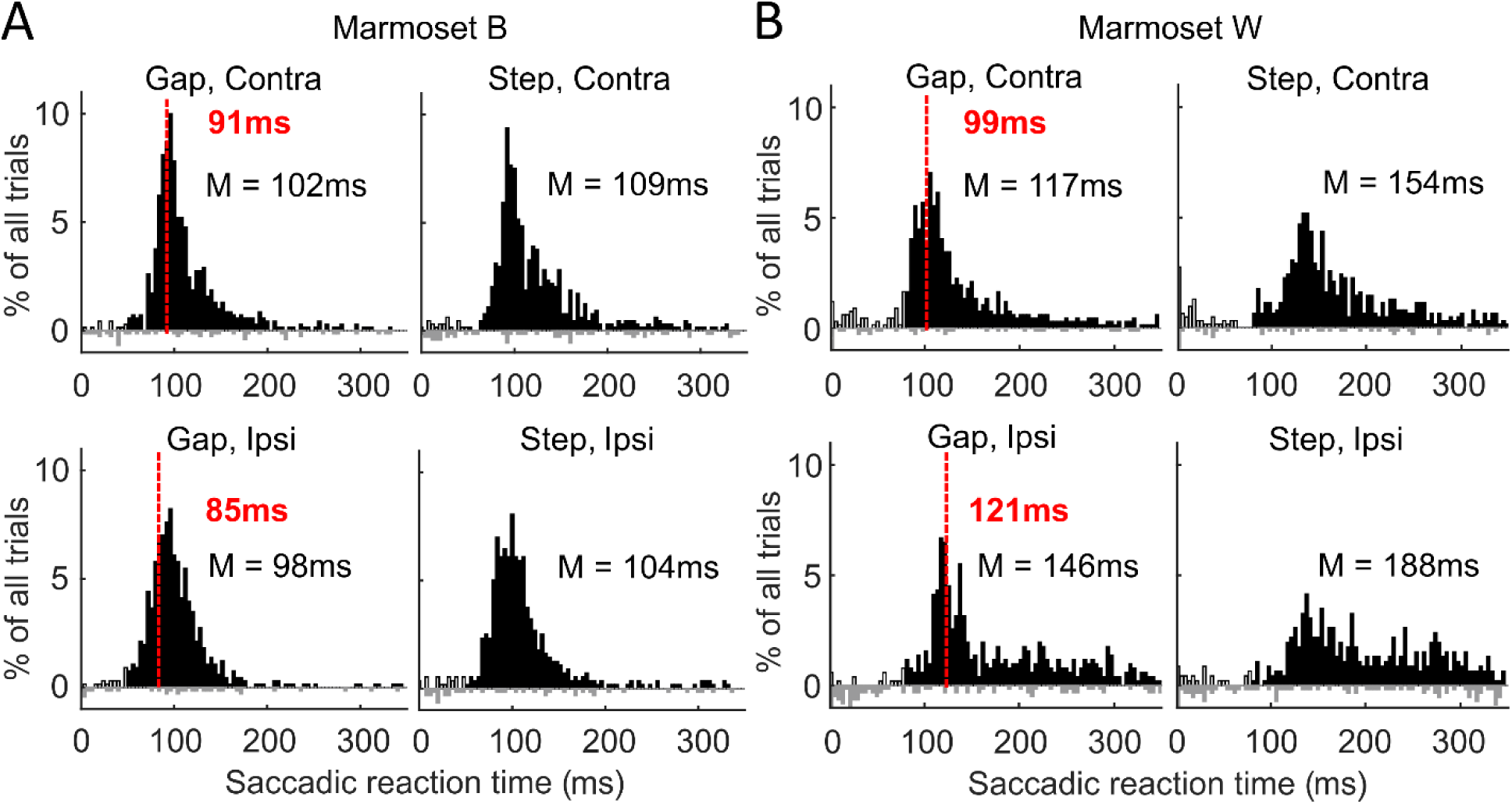
SRT histograms showing the percentage of trials with SRTs in 4-ms bins between 0 and 350ms. Black bars: correct trials; gray bars: error trials; empty bars: anticipatory saccades. Red dashed line: boundary of the shortest quartile of SRTs. Correct saccades with SRTs shown to the left of the line were classified as “fast saccades” and those to the right were “slow saccades”. The median (M) of SRTs was also shown for each trial type and direction. A). SRT histograms of saccades performed by Marmoset B. B). SRT histograms of saccades performed by Marmoset W. SRT: saccadic reaction time, Contra: contralateral saccades, Ipsi: ipsilateral saccades.

### Response of single units to the gap

A total of 361 well-isolated single units were recorded from both marmosets (Marmoset B: n = 173, Marmoset W: n = 188). To identify neurons that responded during the gap, we performed for each unit a paired *t-*test between activity during the peristimulus and fixation periods in Gap trials. The peristimulus period lasted from 165ms before stimulus onset to 34ms after stimulus onset, which in Gap trials was the same as the 200-ms period starting from 35ms after the offset of the fixation point. For the fixation period we chose the 200ms before the offset of the fixation dot, or 400ms to 201ms before stimulus onset. We found 54 cells that responded significantly to the gap (p < 0.05) constituting 15% of the total population and were referred to as the ‘gap neurons’. Their absolute gap-period response, compared with that of the other cells, was illustrated in Figure 4A. We found a significant interaction between cell type and task (mixed model ANOVA, F = 116.0, p < 4.9 × 10^−324^, Figure 4A). While the other cells had comparable responses in either Gap or Step trials (post hoc Tukey’s test: p = 0.61; right bars), the gap cells had significantly greater response during Gap than Step trials (p = 7.7 × 10^−6^; left bars). Not only did they have a greater response during the peristimulus period in Gap trials (p = 7.7 × 10^−6^, filled bars), this was also the case in Step trials (p = 0.0015, empty bars).

**Figure 4.**
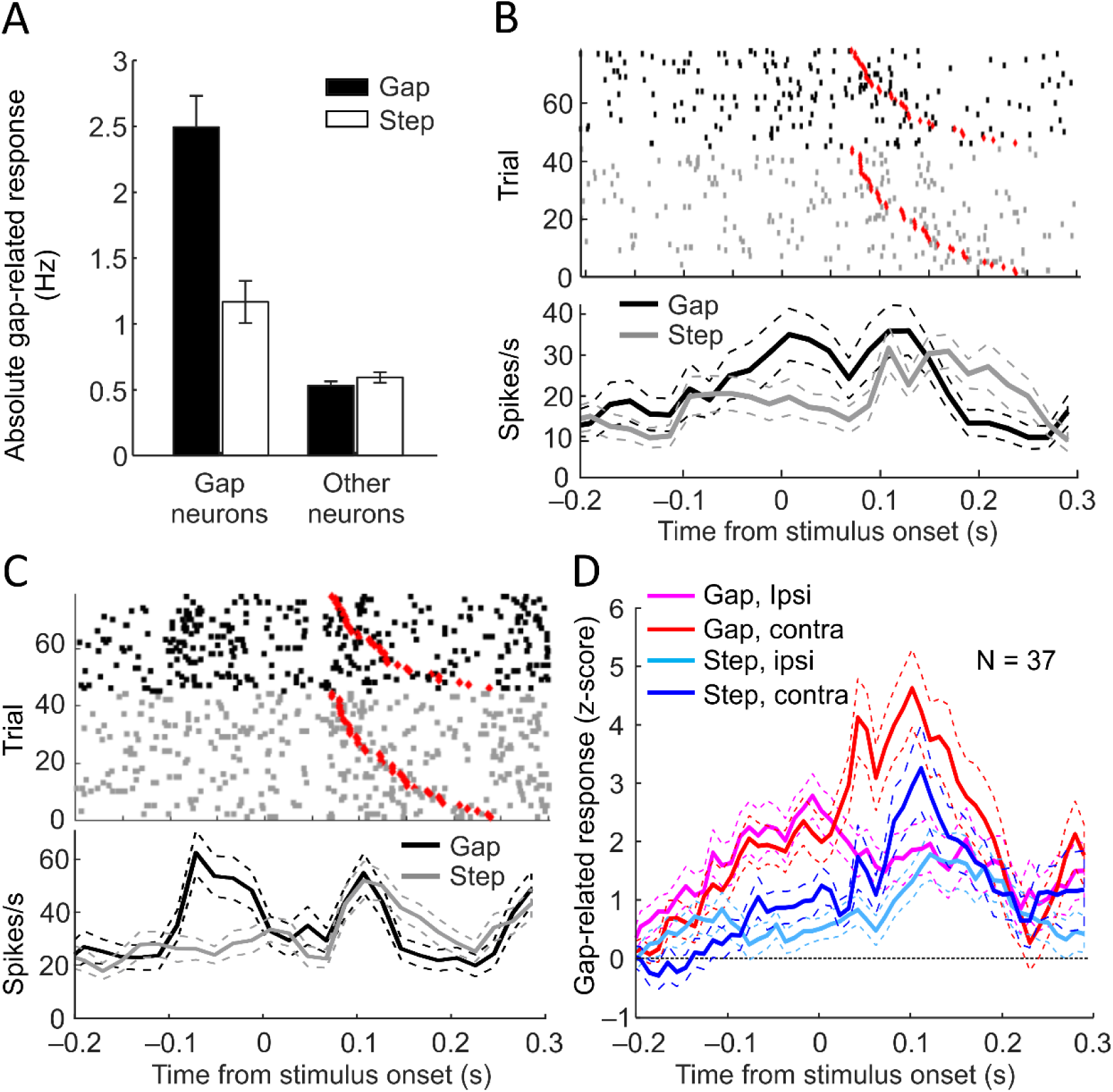
Gap neurons and their peristimulus change in activity. A) Absolute change in firing rates from the fixation to the gap period in gap (left filled bar) versus other (right filled bar) neurons compared with their responses in the peristimulus period in Step trials (empty bars). Gap but not the other neurons responded more strongly in Gap than Step trials. B) Example of a gap neuron which showed gradual increase in activity towards the end of the peristimulus period in Gap but not Step trials. Each tick mark denotes a single action potential. Red diamonds mark saccadic onsets. C). Example of a gap neuron which displayed a sudden increase in activity approximately 100ms following the offset of the fixation dot. Both neurons in B) and C) also showed a target-related response starting approximately 70ms following target onset. D) Standardized peristimulus response averaged across all gap neurons with a positive response. These cells also had a second response to the target, with a preference for contralateral targets emerging at 35ms following their onset.

The activity of representative gap cells is shown in Figure 4B and C. In the raster plots, each black dot represents a single action potential during a Gap trial and a gray dot marks a single spike in a Step trial. Each row illustrates the neuron’s activity from 200ms before to 300ms after stimulus onset in one trial. For each task, the trials were sorted by saccade onset as marked by red diamonds. The curves in both Figure 4B and C display the trial-averaged level of activity in each consecutive 10-ms bins, with the dashed lines depicting the standard error of the mean. Figure 4B shows a neuron with activity that ramped up in the second half of the peristimulus period in Gap (black curve) but not Step trials (gray curve). Figure 4C shows a different type of response among the gap cells: a sharp rise in activity several tens of milliseconds after the offset of the fixation point. Both cells also displayed visual activity approximately 70ms after stimulus onset in both Gap and Step trials. We would characterize such visual responses in the next section.

To visualize the averaged activity of the gap cells, we first z-score standardized the peri-stimulus activity of each neuron against its activity during the fixation period, defined as 400ms to 201ms before stimulus onset. This was done to prevent the high firing-rate neurons activity from dominating the averaged group-level activity. Among the 54 gap neurons, 37 displayed a positive response to the gap period, the averaged activity of which is shown in Figure 4D. During the second half of the gap period—the last 100ms before stimulus onset—and the 35ms after stimulus onset, these neurons had stronger responses in Gap (magenta and red curves) than in Step trials (light and dark blue curves). From 35ms after stimulus onset, the visual signal started to exert an influence on these cells, reflected in sharp increases in averaged activity in trials with contralateral targets (red and dark blue) but not those with ipsilateral targets (magenta and light blue). That is, posterior parietal gap cells in the left hemisphere were visually responsive if the target was presented in the right half of the visual field.

The 2D configuration of the Utah array allowed us to estimate the spatial distribution of the gap cells along the anterior-posterior and medial-lateral axes within the posterior parietal cortex. Figure 5A shows the percentage of the total neuronal population recorded from each recording site. Along the y-axis, the medial edge of the array is shown at the top of the plot, and the anterior edge of the array is to the left of the plot along the x-axis. Channels in the four corners of the arrays together served as ground. Neurons were recorded from most channels, 75.0% and 84.4% respectively for the two marmosets. As expected in random sampling from a smooth cortex, our relative success in sampling neurons at the same sites in the 2D map differed across the two marmosets (Chi-square goodness-of-fit test: χ^2^ = 211.2, df = 31, p = 4.9 × 10^−324^). We then plotted the percentage of gap cells out of all recorded cells at each recording site for each marmoset (Figure 5B). Gap cells were found in 62.5% and 40.7% of all channels from which single units were recorded, respectively for the animals. The distribution of gap cells as a percentage of cells recorded at each site displayed non-uniform, cluster-like topography (Chi-square goodness-of-fit test against uniform discrete distribution for 2 subjects: χ^2^ = 449.8 and 1033.5, df = 23 and 26, p = 4.9 × 10^−324^ for both animals). Given that the arrays straddled the IPS, we asked if gap neurons concentrated more in the lateral half of the arrays which was more likely to be in the LIP according to the atlas. While gap neurons did not concentrate on either medial or lateral half of the array for Marmoset B (unpaired t test, *t*22 = 0.11, p = 0.92), for Marmoset W they were more likely to be detected by the medial half of the channels (*t*25 = 2.42, p = 0.023). We also compared the rate of gap cell detection between the anterior and posterior halves of the arrays and found no difference in either animal (unpaired t test, *t*22 = –0.30 and *t*25 =–1.25, p = 0.77 and 0.22, respectively). The difference in the 2D distribution of recorded cells made it difficult to meaningfully compare the distribution of gap neurons in the PPC across subjects. Despite of this, both marmosets appeared to have a cluster of gap neurons recorded by the medial-posterior area of the arrays (Figure 5B).

**Figure 5.**
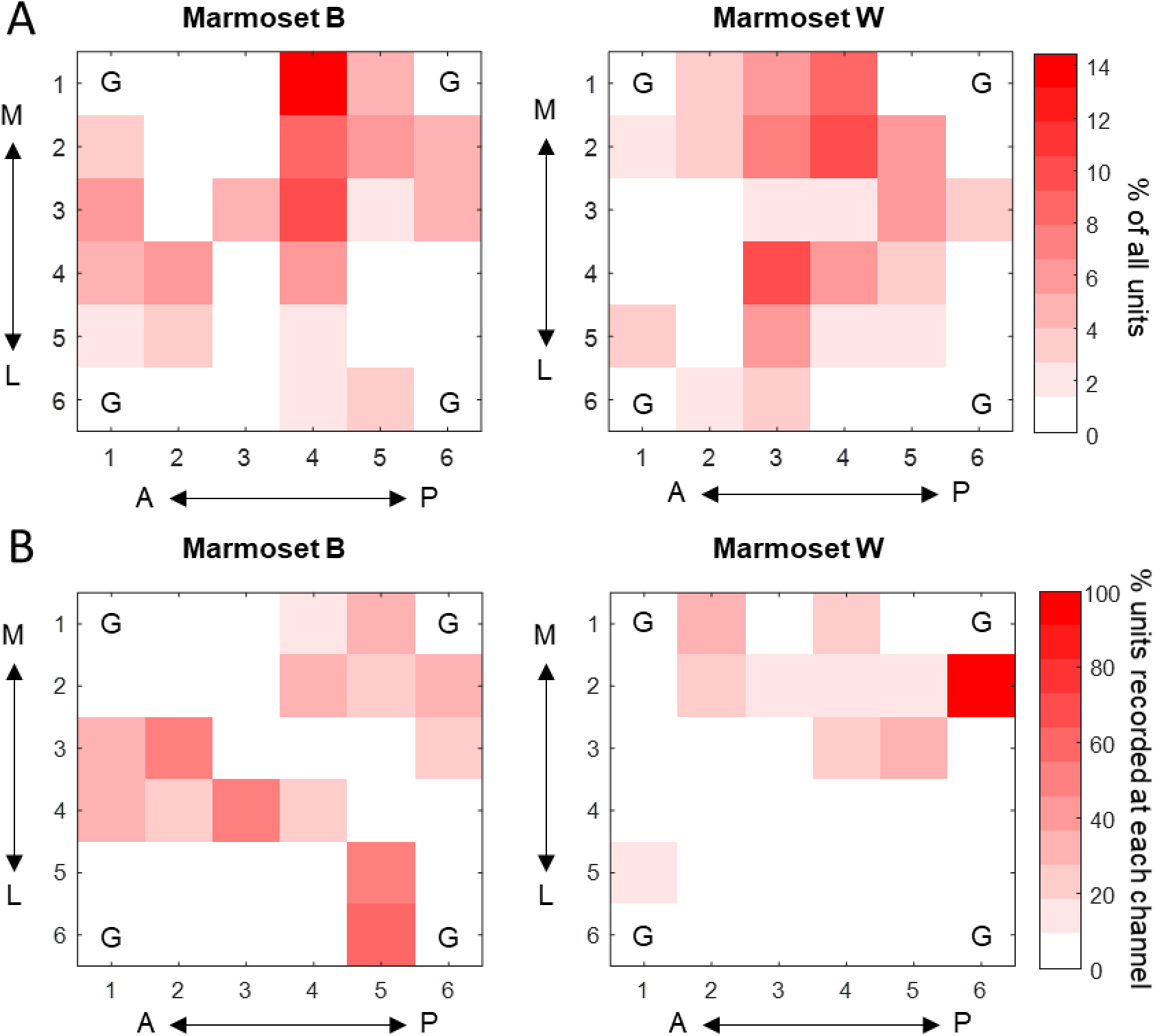
Percentage of all neurons recorded and of gap neurons from each channel as a function of its topographical location in the marmoset PPC. In each panel, the abscissa corresponds to the anterior-posterior axis of the brain, while the ordinate corresponds to the medial (top)-lateral (bottom) axis. A) Percentage of all recorded neurons detected by each recording site in Marmoset B (left) and W (right), respectively. B) Gap neurons as a percentage of all neurons recorded at each site in Marmoset B (left) and W (right), respectively. M: medial, L: lateral, A: anterior, P: posterior, G: grounding channels.

### Response of single units to the saccade target

Because the posterior parietal cortex in macaques is known to contain visually-responsive neurons (Andersen et al., 1985; Colby et al., 1996; Ben Hamed et al., 2001), or ‘target neurons’ for short, we went on to identify neurons that responded to the saccade target. Based on a paired *t-*test between the visual period (35 to 134ms after target onset) and the fixation period (400ms to 201ms before target onset), we identified 142 or 39.3% of all recorded neurons as being significantly responsive to the visual period in at least one type of trials (Figure 6A, left). Notably, the activity of these neurons rose abruptly at 35ms after target onset, suggesting the arrival of the visual signal at the PPC. Among these neurons, 40.9% responded only in Gap trials, 24.7% responded only in Step trials, and the remaining 34.5% responded in both trial types. Together the normalized activity of these 142 target neurons were contrasted with the remaining 219 neurons (60.7%; Figure 6A, right). We conducted a mixed model ANOVA on the neurons’ normalized and averaged response in the peristimulus period, with task (Gap vs. Step) and direction (ipsilateral vs. contralateral) as within-subject variables and cell type as the between-subject factor. We found a significant interaction between cell type and saccadic direction (*F*_1,347_ = 11.4, p = 0.00084): target neurons responded more strongly to contralateral than ipsilateral targets (post hoc Tukey’s test, p = 3.2 × 10^−5^ and 3.3 × 10^−5^ in Gap and Step trials respectively), whereas the other neurons did not show this difference (p = 0.70 and 0.26 in Gap and Step trials). There was also a significant interaction between cell type and the task (*F*_1, 347_ = 6.62, p = 0.011), although the task effect in the response of target neurons had only a trend toward significance (post hoc Tukey’s test, p = 0.086), compared with a lack of difference in the other cells (p = 0.64). We found no interaction between task and saccade direction (*F*_1, 347_ = 1.15, p = 0.28).

**Figure 6.**
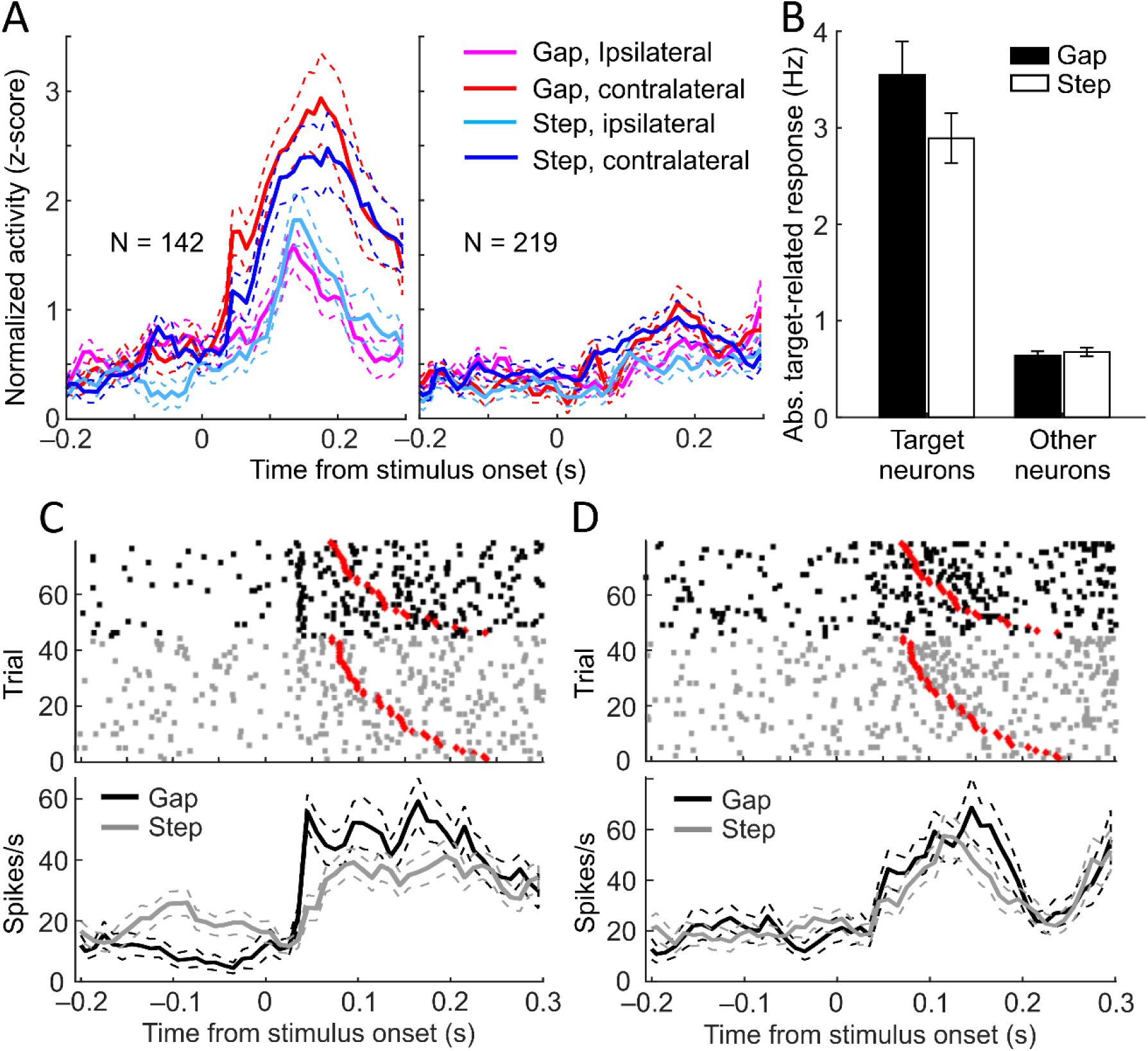
Target neurons and their target-related change in activity. A) Standardized target-related response averaged across all target neurons (left), compared with the response of non-target neurons (right). In target neurons, the response starts at 35ms after target onset, with a strong preference for contralateral targets. B) Absolute change in firing rates from the fixation to the visual period in target (left filled bar) versus other (right filled bar) neurons compared with their responses in the visual period in Step trials (empty bars). In each trial type, the response is stronger in target neurons, which also responded more strongly in Gap than Step trials. C) Example of a gap neuron with a steep rise in activity at approximately 35ms following target onset. This response was more consistently timed in Gap than in Step trials. The same neuron also responded differently during the peristimulus period in Gap and Step trials. Each tick mark denotes a single action potential. Red diamonds mark saccadic onsets. D). Example of a gap neuron which displayed a sudden increase in activity approximately 35ms following target onset. Unlike C), this neuron did not respond to the gap period.

The effect of the gap became significant when the neurons’ *absolute* target-period response was considered. A mixed-model ANOVA with task as within-subject factor and cell type as between-subject factor revealed main effects of both factors, as well as an interactive effect on the neurons’ absolute change in activity from the fixation to the visual period (task: F1,359 = 12.0, p = 0.00059, cell type: F1,359 = 117.8, p < 4.9 × 10^−324^, interaction: F1,359 = 15.3, p = 0.00011; Figure 6B). While target neurons responded significantly to the target in both tasks, their response was stronger in the Gap than in the Step task (post hoc Tukey’s test, p = 2.0 × 10^−5^; Figure 6B, left bars), an effect not observed in neurons that did not respond to the target (p = 0.98; right bars). The target neurons also had stronger response on both Gap and Step tasks than the other neurons (p = 7.7 × 10^−6^ in both tasks). Since the gap and target neurons were identified independently, the two groups were not mutually exclusive. Indeed, 36 neurons belonged to both groups, constituting 10% of all recorded cells, 25.4% of target neurons and 66.7% of gap neurons. Given the existence of these “dual-response” neurons, we asked if they were responsible for the task effect on the target-period activities. To answer this question, we further split the target neurons into dual-response and non-gap target neurons and repeated the mixed model ANOVA with 3 instead of 2 cell types. We found a significant task × cell-type interaction (F2,358 = 12.7, p = 4.9 × 10^−6^): the task effect was contributed solely by dual-response neurons (post hoc Tukey’s test, p = 2.5 × 10^−5^) and not the non-gap target cells (p = 0.10). While these two types of target neurons were similar in their response in Step trials (p = 0.41), the dual-response neurons responded more strongly on Gap trials (p = 0.0032).In short, target neurons preferentially responded to contralateral targets across trial types; 10% of all PPC neurons not only responded to the gap period but also responded more strongly to targets following the gap, potentially contributing to the reduction in SRTs in Gap trials.

Two examples of target neurons are shown in Figure 6C and D, both of which displayed a sharp increase in activity at approximately 35ms following the onset of the target. The cell shown in 6C is also a dual-response cell, displaying an inhibitory response during the gap period and a stronger response to the visual target in Gap trials. Notably, this neuron also encoded a saccade-related signal, as indicated by its consistent reduction of activity after saccadic onset (marked by red diamonds) in Step trials (gray tick marks). Additionally, the neuron shown in 6D also encode an SRT-related signal, as indicated by the sudden increase in its activity following the onset of relatively fast saccades, i.e. in most of the Gap trials and approximately top ½ of all trials shown in the top panel (Figure 6D). Thus, rather than being dedicated to visual signal detection, both neurons were likely involved in visuomotor processing.

We examined the 2D distributions of the target cells as a percentage of recorded neurons from each channel in each of the subjects (Figure 7A). In neither marmoset were the target neurons distributed uniformly across all channels which detected single units (Chi-square goodness-of-fit test: χ^2^ = 290.3 and 785.3, d.f. = 23 and 26 respectively for Marmoset B and W, p <= 4.9 × 10^−324^ for both; Figure 7A). Similar to the case of the gap neurons, this observation suggested a clustered pattern of distribution. To explore whether the target neurons were preferentially distributed either along the medial-lateral or anterior-posterior axis, we compared the rate of their detection between halves of the arrays. We found no difference in target neuron distribution between the medial and lateral halves (unpaired t test, *t*22 = 0.51 and *t*25 = 0.91, p = 0.61 and 0.37, respectively for Marmoset B and W) or between anterior and posterior halves of all channels (*t*22 = 1.30 and *t*25 = 1.00, p = 0.21 and 0.32, respectively). It remains possible that an array with more channels and/or covers a greater area of the PPC can detect a better-defined spatial distribution of target neurons. We then compared the spatial distributions of gap and target neurons, both combined across subjects (Figure 7B). Distinct from Figure 7A, Figure 7B shows the percentage of all gap neurons (left) and target neurons (right) detected from each active channel, hence the numbers in each plot sum to 100%. Overall, these two types of cells did not colocalize (Chi-square goodness-of-fit test: χ^2^ = 50.2, d.f. = 31, p = 0.016). Given the subtle difference in array position across subjects, our finding does not rule out the possibility that the two types of cells were similarly distributed in the PPC.

**Figure 7.**
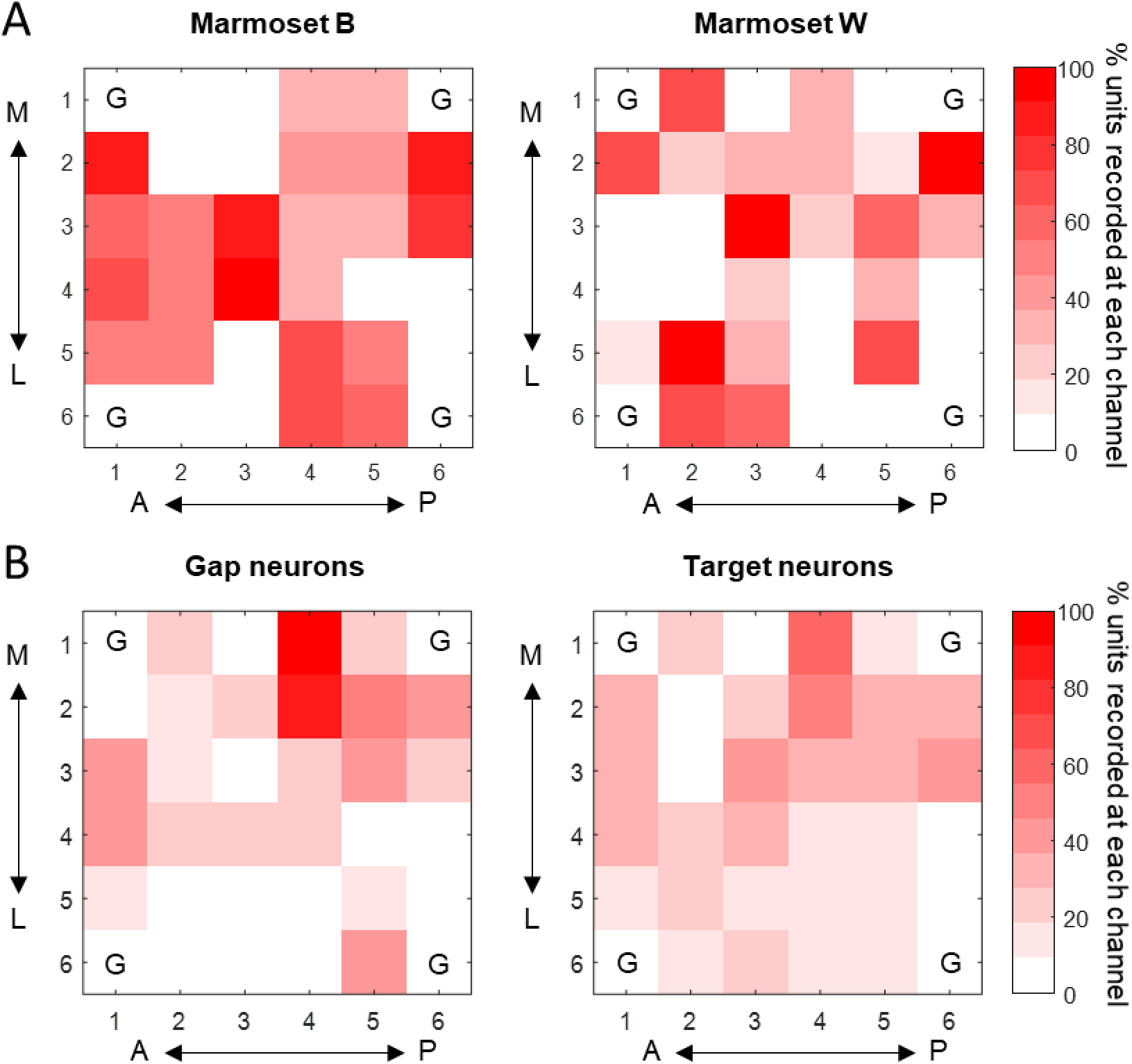
Percentage of target and gap neurons recorded from each channel as a function of its topographical location in the marmoset PPC. In each plot, the abscissa corresponds to the anterior-posterior axis of the brain, while the ordinate corresponds to the medial (top)-lateral (bottom) axis. A) Target neurons as a percentage of all neurons recorded from each channel in Marmoset B (left) and W (right). B) gap (left) and target (right) neurons respectively as a percentage of all neurons recorded at each site, after combining data from both subjects. M: medial, L: lateral, A: anterior, P: posterior, G: grounding channels.

### Correlation between activity change and subsequent saccadic reaction times (SRTs)

Now that we found evidence that neurons in the marmoset PPC responded to the gap and the visual target, we went on to test for the functional relevant of these responses. For relevant groups of neurons, we obtained their peristimulus and/or target-related change in activity in each trial and calculated a correlation coefficient (Spearman’s rho) between this activity change and the subsequent saccadic reaction times (SRTs). If the gap or target cells were part of the circuitry that plans and generates appropriate saccadic responses, then stronger gap or visual responses should precede efficient responses with shorter SRTs.

As a group, the peristimulus response (i.e. change in activity from the fixation to the gap period) of gap cells showed significantly negative correlations with the subsequent SRTs during Gap trials with contralateral targets (one-sample *t* test against 0 with family wise error correction: *t53* = –3.72, padj = 0.0039, black filled bar, left set, Figure 8A). The same was found for Step trials with contralateral targets (*t53* = –2.98, padj = 0.017, gray filled bar, left set) but not for either trial type with ipsilateral targets (padj = 0.44 and 0.59 respectively, empty bars, left set, Figure 8A). Interestingly, cells that did not respond to the gap period significantly (‘Other cells’) also showed negative correlation with the SRTs in contralateral Gap trials (*t303* = –2.44, padj = 0.041, black filled bar, right set, Figure 8A) but not in other trial types (padj >= 0.59). We also performed a mixed model ANOVA to directly compare the correlation coefficients from the two groups of cells, using task and saccade direction as within-subject variables. We found a main effect of cell type (*F*_1,320_ = 8.28, p = 0.0043): gap cells had overall stronger negative correlation than other cells. Thus, these findings support the idea that stronger peristimulus response in gap-sensitive PPC neurons contribute to faster saccadic responses in the Gap task, although their response during the peristimulus period could also contribute to faster saccades in the Step trials. Additionally, this response preparation-related signal was widespread in the PPC and could be detected in neurons that did not respond significantly during the peristimulus period. Intriguingly, such negative correlations were specific to trials with contralateral targets, even though the peristimulus responses took place before the target appeared.

**Figure 8.**
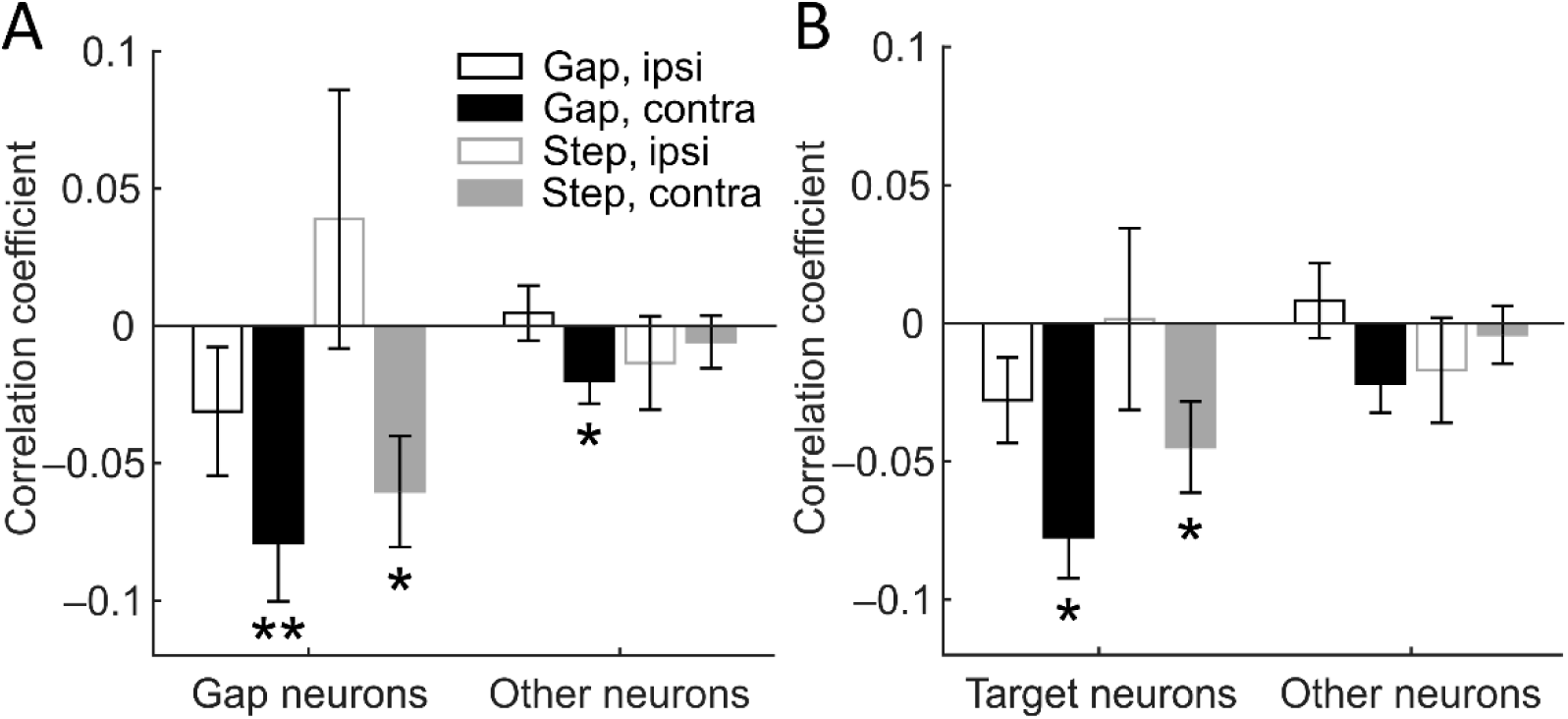
Correlation coefficients (Spearman’s rho) between neuronal response and SRTs. Negative correlations indicate that greater neuronal response preceded shorter SRTs. A) Correlation coefficients between peristimulus-response of gap neurons (left set) and the other neurons (right set) and the SRTs of each type of trials. B) Correlation coefficients between target-related response of target neurons (left set) and the other neurons (right set) and the SRTs of each type of trials. SRT: saccadic reaction time. *p<0.05, **p<0.005

For target neurons, we found that their target-related response (i.e. change in activity from the fixation to the visual period) significantly and negatively correlated with SRTs in contralateral Gap and Step trials (one-sample *t* test against 0 with family wise error correction: Gap: *t141* = –5.18, padj = 6.1 × 10^−6^, black filled bar, Step: *t140* = –2.70, padj = 0.031, gray filled bar, left set, Figure 8B). This was not found for neurons that did not respond during the visual period in either trial types (padj >= 0.11, right set, Figure 8B). Hence the responses of target neurons may also contribute to faster saccadic responses. Spanning 100ms from 35ms after target onset, the ‘visual period’ may also have contained activities related to motor preparation and execution, although the fact that the negative correlations were similar in strength to those found in the peristimulus period (Figure 8A vs. 8B) seems to argue against this idea. Taken together, the findings demonstrate the presence of a saccadic preparation-related signal during both the peristimulus and visual-response periods in the marmoset PPC.

### Changes in gap- or visual-related activity preceding fast vs. slow saccades

As described above (Figure 3), we categorized saccades in Gap trials into ‘fast saccades’, defined as those with SRTs in the shortest quartile; and ‘slow saccades’ which included the rest of the trials. Given that stronger gap- and target-related responses in relevant cell groups preceded shorter SRTs, we expected fast saccades to be preceded by stronger neuronal responses as well.

Since changes in neuronal response could be positive or negative, we analyzed them separately. Given that the relationship between gap-related response and SRTs depended on target location (Figure 8A), we also separately analyzed trials involving contralateral versus ipsilateral saccades. For neurons with positive gap-related responses in trials involving contralateral saccades, a 2-way ANOVA was performed with cell type (gap vs. other) and saccade type (fast vs. slow) as the two factors. We found main effects of both saccade type (*F*_1,333_ = 26.3, p = 4.9 × 10^−7^) and cell type (*F*_1,333_ = 11.1, p = 0.00095; Figure 9A, top left panel). A post hoc Tukey’s test revealed significant greater gap-related response preceding fast saccades in both gap cells (p = 0.024, left bars) and in other cells as well (p = 7.8 × 10^−6^, right bars, Figure 9A, top left panel). In trials with ipsilateral targets, the same analysis revealed quite a different pattern (Figure 9A, top right panel). We found no effect of cell type (*F*_1,315_ = 2.78, p = 0.096) or saccade type (*F*_1,315_ = 0.013, p = 0.91). In short, fast saccades to contralateral targets were preceded by greater gap-related activity in PPC neurons.

**Figure 9.**
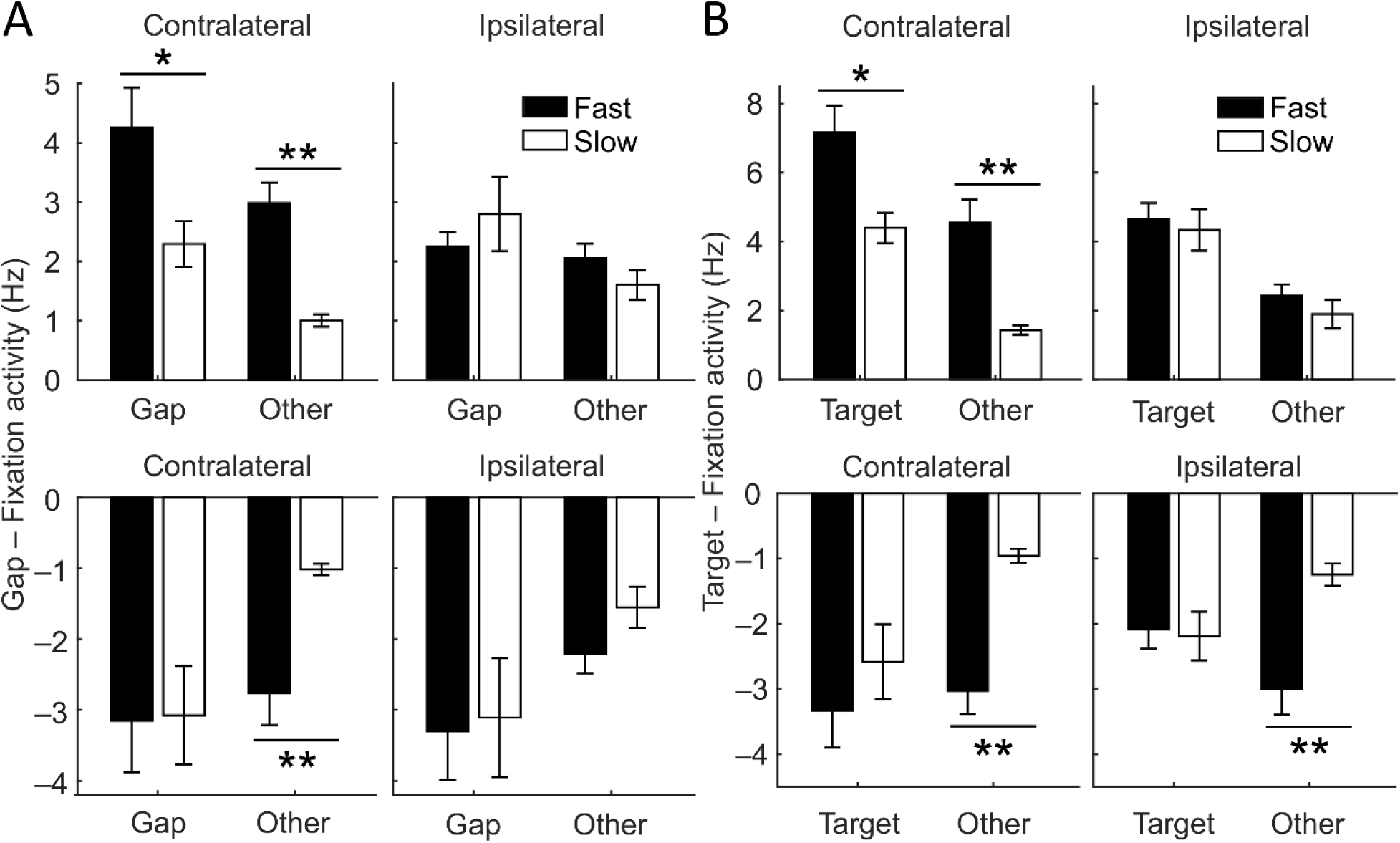
Neuronal response during the gap and visual periods in Gap trials with fast and slow SRTs. Fast SRTs were defined as those in the shortest quartile of the SRT distribution. A) In trials with contralateral targets, gap neurons with a positive response (i.e. those with a significant increase in firing rate in the gap period) did so more strongly in the gap period before fast than slow saccades (left bars, top left). The other neurons with a non-significant increase in activity also did so more strongly before fast than slow saccades (right bars, top left). Gap neurons with an inhibitory response to the gap period did not respond more strongly before fast saccades (left bars, bottom left), although the remaining neurons had a greater reduction in activity before fast saccades (right bars, bottom left). In trials with ipsilateral targets, neither type of neurons responded differently before fast and slow saccades (right panels). B) In trials with contralateral targets, target neurons with a positive response (i.e. those with a significant increase in firing rate in the visual period) responded more strongly in the visual period before fast than slow saccades (left bars, top left). The other neurons with a non-significant increase in activity also did so more strongly before fast than slow saccades (right bars, top left). Target neurons with an inhibitory response to the gap period did not respond more strongly before fast saccades (left bars, bottom left), although the remaining neurons had a greater reduction in activity before fast saccades (right bars, bottom left). In trials with ipsilateral targets, while neurons with positive responses—significant or not—did not respond differently before fast and slow saccades (top right), non-target neurons showed a greater reduction in visual-period activity before fast than slow saccades. SRT: saccadic reaction time. *p<0.05, **p<0.0005

We then examined the cells that displayed a negative gap-related response, i.e. decreased in activity level from the fixation to the gap period. In contralateral-saccade trials, we found an effect of cell type (*F*_1,258_ = 4.99, p = 0.026) but not of saccade type (*F*_1,258_ = 2.78, p = 0.097) or any interaction between the two (*F*_1,258_ = 2.32, p = 0.13, Figure 9A, lower left panel). Interestingly, we found a stronger response preceding fast saccades in the other cells (post hoc Tukey’s test, p = 0.00025) but not in gap cells (p > 0.99). Thus, while the gap neurons had generally greater (negative) changes in activity compared to the other cells, such response did not precede fast saccades. In trials with ipsilateral targets, we found an overall effect of cell type (*F*_1,230_ = 5.27, p = 0.022) but not of saccade type (*F*_1,230_ = 0.55, p = 0.46) or any interaction between the two (*F*_1,230_ = 0.17, p = 0.68, Figure 9A, lower right panel). Taken together, while stronger gap-related responses were more likely followed by fast saccades, this was only the case when the targets were presented contralateral to the PPC neurons.

Figure 8B demonstrated a strong relationship between the target response of PPC neurons and the SRTs, in a manner also dependent on the target location. We therefore conducted a similar set of 2-way ANOVAs for the target responses. On positive responses, we found effects of both saccade type (*F*_1,393_ = 33.1, p = 1.7 × 10^−8^) and cell type (*F*_1,393_ = 29.6, p = 9.2 × 10^−8^) in contralateral-saccade trials (Figure 9B, top left panel). Both target cells and other cells had significantly greater positive visual response before fast saccades than before slow saccades (post hoc Tukey’s test, p = 0.0015 and 0.00039, respectively). By contrast, among trials with ipsilateral targets, we did not find any effect of saccade type (*F*_1,309_ = 0.74, p = 0.39), although target neurons still had stronger response than other cells across saccade types (*F*_1,309_ = 21.9, p = 4.3 × 10^−6^). For cells with negative responses, in trials with contralateral targets, we found both significant effects of saccade type (*F*_1,209_ = 15.4, p = 0.00012) and of cell type (*F*_1,209_ = 7.2, p = 0.0079), which were explained by a greater target-related response in other cells in fast-saccade trials (p = 1.3 × 10^−5^) but not in target cells (p = 0.59). In trials with ipsilateral targets we also found effects of saccade type (*F*_1,246_ = 5.56, p = 0.019) and of an interaction between the factors (*F*_1,246_ = 7.08, p = 0.0083), also explained by a greater target-related response in other cells in fast-saccade trials (p = 8.8 × 10^−5^) but not in target cells (p > 0.99). Taken together, similar to the case of gap-related responses, fast saccades, especially those directed at contralateral targets, were associated with stronger visual responses in the entire recorded population of PPC neurons.

## DISCUSSION

The common marmoset is a promising primate model for human cognition and social interaction. Since saccades provide an essential tool for quantifying complex cognitive processes, it is essential to obtain a detailed understanding of the cortical mechanisms of saccadic control in the marmoset. Here we recorded single unit activity through chronically implanted microelectrode arrays in the PPC while marmosets performed visually guided saccades with or without a gap between fixation offset and target onset. Marmosets demonstrated a gap effect similar to humans and macaques (Johnston et al., 2018). We found that 15% of all PPC units recorded responded significantly to the gap, and their response magnitude negatively correlated with subsequent SRTs on trials with contralateral targets. The remaining 85% of PPC neurons on averaged also had a small but significant negative correlation with SRTs on contralateral trials. Additionally, we found 39% of PPC units that responded to the peripheral target, and greater responses in them also preceded shorter SRTs on Gap trials and contralateral Step trials. Both types of cells showed stronger response before express than regular saccades. Importantly, in *the population of* PPC cells recorded, both the gap-related and target-related responses were stronger before fast saccades than slow saccades to contralateral targets, which strongly support a role of the marmoset PPC in modulating saccadic preparation. Our findings suggest the presence of an area homologous to the macaque LIP in the marmoset PPC. Existing literature and the current study together suggest that the PPC contains a key oculomotor area across primate species. In human patients with posterior parietal lesions, while the deficits in spatial attention may be the most striking, their impairment in saccadic performance is no less severe (Ptak and Müri, 2013). These patients have markedly increased latencies especially for contralateral saccades (Pierrot-Deseilligny et al., 1991; Braun et al., 1992) and significant reduction in express saccades in a gap paradigm (Braun et al., 1992). Our finding of gap-responsive cells in the PPC was consistent with a role of the area in oculomotor functions in the marmoset, similar to that of the LIP in macaques and parietal eye field in humans.

To establish homology in cortical areas across species, three lines of evidence need to be considered: cytoarchitecture, connectivity, and neural response properties (Kaas, 1987; Krubitzer, 1995). In the marmoset brain atlas by Paxinos et al. (2012), the PPC is parcellated based on cyto- and myelo-architecture, using the IPS as a reference. Specifically, Rosa et al. (2009) identified two subdivisions in the marmoset PPC based on the pattern of myelination and soma size of layer V pyramidal neurons in a way consistent with the parcellation in the macaque PPC (Blatt et al., 1990). They further suggested the LIP to be the subarea with the heaviest myelination within the dorsal subdivision (Rosa et al., 2009). According to the atlas (Paxinos et al., 2012), since our arrays straddled the intraparietal sulcus (IPS) as revealed by ex vivo MRI and in vivo micro-CT scan, it would have covered part of the MIP, VIP and LIP, located in the medial and lateral banks and the fundus of the IPS, respectively. Also, our 2.4 × 2.4mm array should have resided within the anterior and posterior borders of the LIP, which extends 3.5-4mm rostro-caudally (Paxinos et al., 2012). By contrast, some have suggested the marmoset LIP to be an area that includes both banks of the IPS, based on the presence of calbindin-rich deep layers (Bourne et al., 2007). In addition to cytoarchitecture, functional connectivity and tracing studies have supported the presence of an LIP homologue in the marmoset PPC, given its connections with the FEF and the SC (Reser et al., 2013; Majka et al., 2016; Ghahremani et al., 2017). In macaque monkeys, the LIP stands out as the only parietal region that projects directly to the SC (Lynch et al., 1985; Andersen et al., 1990). In comparison, in the marmoset PPC, the corticotectal neurons appear more distributed, and their locations among the atlas-defined subareas remain unclear (Collins et al., 2005). In macaque monkeys, reciprocal connections exist between the FEF and the LIP and VIP, but not MIP (Stanton et al., 1995, 2005). By contrast, in the marmoset, all three areas as defined by the atlas are reciprocally connected with area 8aV (Reser et al., 2013; Majka et al., 2016), the putative marmoset FEF. Thus, the existence of an area homologous to the macaque LIP in the marmoset PPC is supported by studies of cytoarchitecture and connectivity, but its precise boundary remains unclear and appears to extend beyond the lateral bank of the IPS.

The current study is one of the first to provide evidence concerning the third criterion for homology—neural response properties (Kaas, 1987; Krubitzer, 1995). In macaque monkeys, distinct from the LIP, the MIP is specialized for reaches (Gnadt and Andersen, 1988; Snyder et al., 1997) and its lesion does not affect contralateral saccade choices (Christopoulos et al., 2015). If the functional differentiation in the marmoset PPC followed the same pattern as in macaques, gap-sensitive neurons should be detected more frequently on the lateral side of the array, which was not the case. Consistent with this finding, microstimulations through the same arrays implanted in the same marmosets triggered no bodily movement beyond saccades and eye blinks, and the likelihood and amplitude of evoked saccade did not increase from the medial to the lateral side of the arrays (Ghahremani et al., 2019). In the macaque PPC, it is now clear that oculomotor and visuomanual signals are mixed within areas either primarily involved in eye (Dickinson et al., 2003) or arm movements (Battaglia-Mayer et al., 2001; Archambault et al., 2009). While the evidence suggests cross-modal integration and gradual transitions from one functional domain to another in the macaque PPC (Hadjidimitrakis et al., 2019), in the marmoset PPC no such gradual change in neuronal activity or evoked response (Ghahremani et al., 2019) were observed. Taken together, the three lines of evidence supports the existence of a marmoset homologue of the macaque LIP in the PPC and suggest that this homologue may be located differently with reference to the IPS.

The specific oculomotor function played by the primate PPC remains a topic of active investigation. Rather than being directly involved in saccade planning, the PPC likely contribute to the modulation of saccade through its role in attention and perception (Bisley and Goldberg, 2003). Previous studies have identified visual neurons (Andersen et al., 1985), e.g. those responding to the onset (Gottlieb et al., 1998; Kubanek et al., 2013) or offset (Ben Hamed and Duhamel, 2002) of salient or relevant targets; as well as neurons responsive to both visual target and saccadic onset (Colby et al., 1996; Gottlieb et al., 2005), with differential distribution along the dorsal-ventral extent of the macaque LIP (Chen et al., 2016). Similarly, in the marmoset PPC, we observed visual neurons responsive to fixation offset and those responsive to target onset and saccadic onset. We speculate that the gap-period activities in the PPC can directly enhance the pre-target activities observed in SC saccade neurons in macaques (Dorris et al., 1997), and those responsive to the target can provide additional input that drives SC neurons over the saccade threshold. For fast saccades, the gap-period input from the PPC may be more important than the target-related input, as the short timing of these saccades suggests a direct pathway that relays the target information from the visual cortex to the brainstem saccade generators via the SC (Fischer, 1987; Schiller et al., 1987; Matsue et al., 1994; Edelman and Keller, 1996; Dorris et al., 1997). In the macaque FEF, it was found that the neurons showing a gap-related activity decrease preferentially projected to SC fixation neurons (Sommer and Wurtz, 2000). A decrease in the activity of these FEF neurons can therefore weaken the excitatory input to SC fixation neurons, thereby disinhibit SC saccade neurons. In the marmoset PPC, we found neurons that displayed greater reduction in activity before fast saccades and speculate that they may similarly contribute to reduced excitation of fixation neurons and disinhibition of saccade neurons in the SC. Together these observations can help explain the significant reduction in express saccades following posterior parietal lesion (Braun et al., 1992).

Although the percentage of gap-sensitive neurons found here may appear less than in macaque LIP (Chen et al., 2013), it should be noted that we included all single units detected in our chronic arrays without imposing any criterion other than a firing rate of greater than 0.3Hz. Strikingly, the rest of the neuronal population, despite not having a significant gap-related response, still responded more strongly during the gap period before fast than slow saccades to contralateral targets. This demonstrates the prevalence of the facilitatory signal for contralateral motor preparation that preceded the target. The lack of significance in the majority of PPC neurons, due to either a smaller magnitude or lack of consistency across trials in their gap-related response, is not unexpected in an association area where individual neurons often respond to multiple variables and reliable representation is achieved at the neuronal ensemble level (Zhang et al., 2017). Overall, during the gap period, both the subtle but widespread signal and the strong response in a subset of PPC neurons likely contributed to the gap effect and express saccade generation by increasing the excitation in saccade neurons in the SC (Everling and Munoz, 2000).

Notably, the gap-related response of the entire recorded neuronal population negatively correlated with the subsequent SRTs on Gap trials with contralateral targets, although the correlations were stronger in gap-sensitive neurons. In macaque monkeys, this phenomenon has been observed in the LIP (Chen et al., 2013), as well as in the SC (Dorris et al., 1997; Dorris and Munoz, 1998; Everling et al., 1999) and the FEF (Everling and Munoz, 2000), which suggests a concerted preparatory process at the circuit level. Our finding indicates that this circuitry and process are shared across the two species—a hypothesis that will need to be tested by single unit recordings from other oculomotor areas in the marmoset. It should be noted that the gap-period activity of PPC neurons only negatively correlated with the SRTs when the targets were contralateral, even though their gap-related response was independent of the target location. These neurons also went on to show a second response that was stronger following the onset of contralateral targets. Both observations suggest that their gap-related response only contributed to the preparation for saccades to contralateral targets, likely via the ipsilateral PPC-SC pathway which controls saccades contralaterally. This directionality also existed for the difference between fast and slow saccades. Together these findings support a role of the PPC in advanced motor preparation for the gap effect and express-like saccades (Munoz and Fecteau, 2002; Chen et al., 2013, 2016). They are not consistent with the effect of enhanced covert attention following fixation disengagement (Fischer, 1987; Braun and Breitmeyer, 1988; Mackeben and Nakayama, 1993), which would predict a correlation between neural responses and SRTs of saccades in both directions. Meanwhile, it is known that the generation of express saccades requires the target and its location to be somewhat predictable, reflecting a component of learning and memory (Klein, 1980; Kalesnykas and Hallett, 1987; Becker, 1989; Kowler, 1990; Paré and Munoz, 1996; Schiller et al., 2004).

In target cells, we also found that their targeted-related activity was correlated with subsequent SRTs, and their target-related activity was stronger before fast than slow saccades. LIP visual neurons defined by a transient response to the target in a memory-guided saccade task did not show activities correlated with the SRTs in the gap task (Chen et al., 2013). Since our marmosets did not perform memory-guided saccades on the same sessions analyzed here, we were unable to isolate the purely visual neurons that may exist in the marmoset PPC and did not intend to liken the target cells to the LIP visual neurons. In fact, as shown in the Results section, 2/3 of the gap cells were also target cells, constituting the 25% of target cells that also showed a stronger target-related response in Gap trials. Additionally, the activity of target cells showed a strong effect of target location/saccade direction. Given the 100-ms interval used in the definition of the target cells, these neurons may also be involved in modulating the preparatory signal in the oculomotor circuitry and contribute to the gap effect in general via projections to the SC.

In summary, we conducted the first single-unit recording study in the marmoset PPC and observed both peristimulus and target-related activities, which were correlated with the SRTs especially on contralateral Gap trials. The neuronal population including all units detected responded more strongly during the gap period preceding fast than slow saccades, consistent with the critical role of the PPC in express saccades in humans (Braun et al., 1992). Together our findings support the hypothesis that the marmoset shares a homologous oculomotor network with the macaque monkeys, as suggested by a functional connectivity study (Ghahremani et al., 2017). We suggest that the common marmoset is a highly valuable model for understanding the circuit-level dynamics underlying oculomotor processes and the pathological changes that produce the well-documented oculomotor deficits observed in neuropsychiatric disorders.

## Acknowledgements

This research was supported by a Foundation grant from the Canadian Institutes of Health Research (FRN 148365) to S.E. and postdoctoral fellowships from the Canadian Institutes of Health Research and the BrainsCAN initiative to L.M.

